# EXPRESSION OF ANO1 IN HUMAN GASTROINTESTINAL TRACT DURING EMBRYONIC AND FETAL DEVELOPMENT

**DOI:** 10.64898/2025.12.30.697055

**Authors:** Vladimir Petrović, Aleksandra Veličkov, Marko Jović, Julija Radenković, Braca Kundalić, Dušan Miljković, Vukota Radovanović, Goran Radenković

## Abstract

Anoctamin 1 (ANO1, TMEM16A) is a transmembrane protein belonging to the ANO family, with a role in the formation of calcium-activated chloride channels (CaCC). It is included in the regulation of physiological processes such as muscle contraction, gastrointestinal motility, secretion, and electrical excitability. Also, recent data suggest that ANO1 is a specific marker for interstitial cells of Cajal (ICC). The aim of the paper was to examine the spatial and temporal distribution of ANO1 in the stomach, small intestine, and large intestine during embryofetal development as a potential marker for the differentiation of ICC and smooth muscle cells. As a material, we used 2 human embryos and samples from 21 human fetuses. The tissue samples were routinely processed into paraffin blocks, and 5 µm-thick sections were immunostained for ANO1. Our results show that ANO1 appears during embryonic development (8^th^ week), and its expression continues through the fetal stages. Epithelial, endothelial and ICC cells consistently expressed ANO1 in all examined samples. Smooth muscle cells showed strong expression in muscularis propria, however by the 25^th^ week this immunopositivity was absent from outer muscle layers in stomach and large intestine. In conclusion, ANO1 can be considered as a reliable marker for following the differentiation of SMC and ICC during embryonic and fetal development.

## Introduction

Anoctamin 1 (ANO1, TMEM16A) is a plasmalemmal protein with eight transmembrane domains, belonging to the ANO family (ANO1-10). It has a role in the formation of calcium-activated chloride channels (CaCC) that facilitate the passive transport of chloride ions into the cytoplasm, and is therefore included in the regulation of physiological processes such as muscle contraction, gastrointestinal motility, exocrine and endocrine secretion, and electrical excitability (1,2). Expression of ANO1 was found in epithelial cells in many organs (salivary glands, lacrimal gland, exocrine pancreas, bronchial tree, intestines, choroid plexus, retina), some types of smooth muscle cells, interstitial cells of Cajal, vascular endothelial cells, and myocardium (3,4). It has been reported that abnormal ANO1 expression might be associated with the pathogenesis of diseases such as cystic fibrosis, hypertension, and gastrointestinal motility disorders (3,5). Besides this, ANO1 overexpression has been observed in many cancers, where it promotes tumorigenesis by influencing cancer cell proliferation, survival, and migration (6-9).

ANO1 has recently gained attention as the marker of interstitial cells of Cajal (ICC) (10,11). Interstitial cells of Cajal, originating from c-kit-positive mesenchymal precursors in the primitive gut, are observed at the end of embryonic development (12,13). They first appear in the oesophagus and stomach, and then, following the rostrocaudal pattern of development, in the small and large intestine. Their development is closely associated with the colonisation of the digestive tube by neural crest cells, which will eventually give rise to the neurons and glial cells of the myenteric and submucosal plexuses (14). ICC can first be distinguished by c-kit immunopositivity at the end of embryonic development, and by the 11th week, they completely surround the ganglia of the myenteric plexus (14-16).

The ICC are regarded as a crucial component of the enteric nervous system, providing the physiological basis for peristaltic movements (17,18). In response to cholinergic stimulation, ICC generate slow, non-oscillatory intestinal contractions that depolarise the ICC-smooth muscle cell network and convert the excitatory message from the motoneurons into muscle contractions (1, 19). The spontaneous pacemaker activity generated by ICC, which is conducted to smooth muscle cells, enables electrical slow waves and phasic contraction. In addition, ICC serve as stretch receptors and participate in the reflex peristalsis pathway due to the stretching of the digestive tube by the food content (20). Available data suggest that ANO1 is expressed in all ICC classes, even those that do not contribute to slow-wave generation, implying that ANO1 may have an alternate function in these cells (21). The lack of differentiation and the absence of ICC lies in the pathogenesis of many motility disorders in the gastrointestinal tract. Furthermore, mice lacking ANO1 were reported to have fewer proliferating ICC in culture, suggesting that ANO1 may also be involved in ICC proliferation (11,22). Lower or absent ANO1 expression has been reported in patients with gastrointestinal disorders, including diabetic gastroparesis, suggesting that this protein may play a role in the pathogenesis of these conditions (23,24).

Considering the limited data available in the literature regarding the pattern of ANO1 expression in the human gut tube during development, the aim of this paper was to examine the spatial and temporal distribution of ANO1 in the stomach, small intestine, and large intestine during embryofetal development as a potential marker for the differentiation of ICC and smooth muscle cells, that is, the separation of these two cell populations from the common precursors.

## Material and methods

### Material

The study material comprised 2 human embryos and 21 human fetuses, with gestational age ranging from 8 to 25 weeks (Table 1). The specimens were obtained from the Centre for Pathology (University Clinical Centre Niš, Serbia) after legal abortions and premature births due to intrauterine fetal deaths, according to the principles of the Ethical Committee. Both sexes are represented in the sample, and no specimens had gastrointestinal disorders. Gestational ages were estimated by anatomical criteria according to the Carnegie Staging system, and by crown-rump length, head circumference, and foot length.

**Table 1.** The summary of the material used for study, presented by the number of tissue samples, period of development and gestational age in weeks.

| Number of samples | Period of development | Gestational age (in weeks) |
| --- | --- | --- |
| 2 | Embryo | 8 |
| 1 | Fetus | 10 |
| 2 |  | 11 |
| 1 |  | 14 |
| 3 |  | 15 |
| 3 |  | 17 |
| 3 |  | 19 |
| 4 |  | 22 |
| 4 |  | 25 |

Each specimen was fixed in 10% neutral formalin for 24h and routinely processed into paraffin blocks. 4µm tissue sections were cut on a microtome, adhered to slides, and stained with hematoxylin and eosin and immunohistochemically. Haematoxylin and eosin staining was used to confirm the normal morphology of all samples, consistent with their gestational age.

### Immunohistochemistry

After deparaffinization in a thermostat and xylene, the tissue slides were rehydrated in decreasing concentrations of ethanol (100% and 96%) and distilled water. After heat-induced antigen retrieval (30 minutes), the tissue peroxidases were blocked with a 3% hydrogen peroxide solution for 10 minutes. The antibodies were applied overnight at 4°C (Table 2) and staining was continued the following day by using a secondary antibody conjugated with horseradish peroxidase for 30 minutes (Real EnVision System for visualization (Dako, catalogue number: K5007). Between the steps, the tissue slides were rinsed in TWEEN buffer (pH=7.4). Diaminobenzidine (DAB) was used as the chromogen, after which the slides were dehydrated through a series of increasing ethanol concentrations (96%, 100%), cleared in xylene, and mounted using Canada balsam and cover slips.

**Table 2.** The antibodies used for immunohistochemical analysis.

| Antibody | Catalog number | Supplier | Dilution | Target cell |
| --- | --- | --- | --- | --- |
| Rabbit polyclonal antibody against ANO1 | ab53212 | Abcam | 1:50 | Cells expressing ANO1 |
| Mouse monoclonal antibody against NSE | M0873 | Dako | 1:100 | Ganglionic cells of the myenteric plexus |
| Rabbit monoclonal antibody against desmin | ab32362 | Abcam | 1:300 | Smooth muscle cells in gastrointestinal tract |

### Descriptive analysis

Three sections were analysed from each Microscopic slides were analysed using the Olympus BX50 light microscope (Olympus, Japan) equipped with a digital camera, Leica DFC295 (Leica Microsystems, Germany), at the Department of Histology and Embryology, Faculty of Medicine, University of Niš.

## Results

### Embryonic development

Our results show that ANO1 expression can be observed in the stomach, small and large intestine at the 8th week of embryonic development (Fig. 1A).

**Figure 1.**
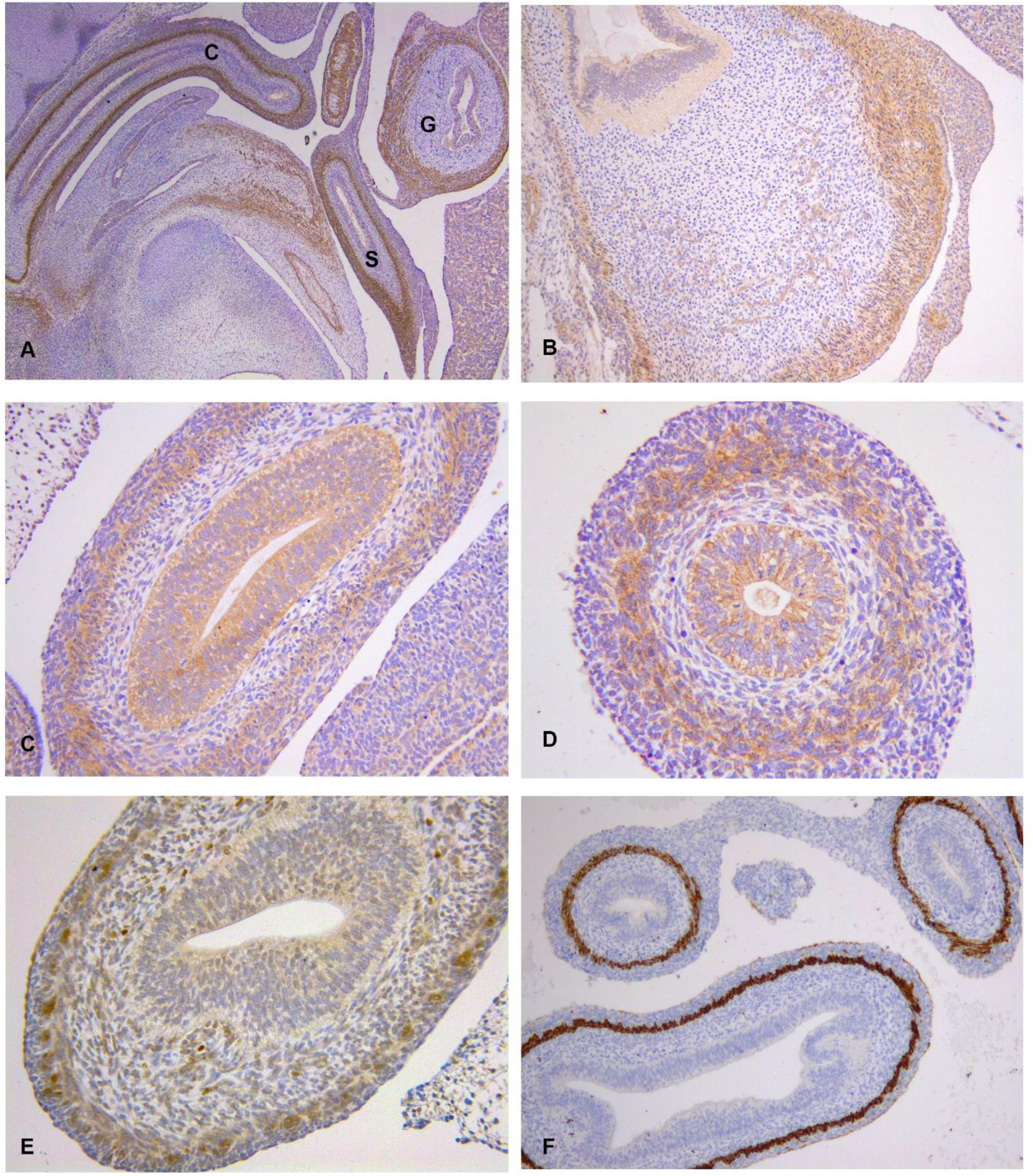
A) Panoramic view of ANO1-immunopositivity in small and large intestine in late embryonic period, 8th week, x40, G-stomach, S-small intestine, C-large intestine; B) ANO1 immunopositivity in the stomach in 8th week of development. The ANO1-positivity is seen in muscular layer, endothelial cells of small blood vessels in submucosa and epithelial cells, x100; C) ANO1 immunopositivity in inner muscular layer and epithelial cells in small intestine, 8th week of development, x200; D) ANO1 immunopositivity in inner muscle layer and epithelial cells in large intestine, 8th week, x200; E) NSE-immunopositivity in ganglionic cells of myenteric plexus in small intestine in 10th week, x200; F) Desmin immunopositivity in smooth muscle cells in small and large intestine in 8th week, x125.

The muscularis propria of the stomach included both the broader inner circular layer and the thinner, outer longitudinal muscle layers that showed strong positivity to desmin. The outer longitudinal layer was observed surrounding the inner layer throughout the entire circumference and consisted of 1-3 rows of cells (Fig. 2F). Developing myenteric plexuses, containing NSE-positive cells, were identified between the two muscle layers (Fig.2D). ANO1-immunopositive cells were observed in both the inner and outer layers of muscularis propria, whereas the large oval ganglionic cells in the myenteric plexuses were ANO1-immunonegative (Fig. 1B). In the small and large intestines, only the inner muscular layer was present, containing ANO1-positive cells, while myenteric plexuses were absent (Fig. 1C, 1D, 1F). The ANO1-positive cells in the developing muscularis propria across all examined parts of the digestive tube formed a continuous layer and exhibited a pleomorphic appearance with euchromatic nuclei. Some appeared round, lacking cytoplasmic processes, while others adopted a spindle-shaped phenotype. ANO1-positivity was also observed in the endothelial cells of developing blood vessels in the submucosa, as well as in epithelial cells of the pseudostratified epithelium in the stomach, small and large intestine.

**Figure 2.**
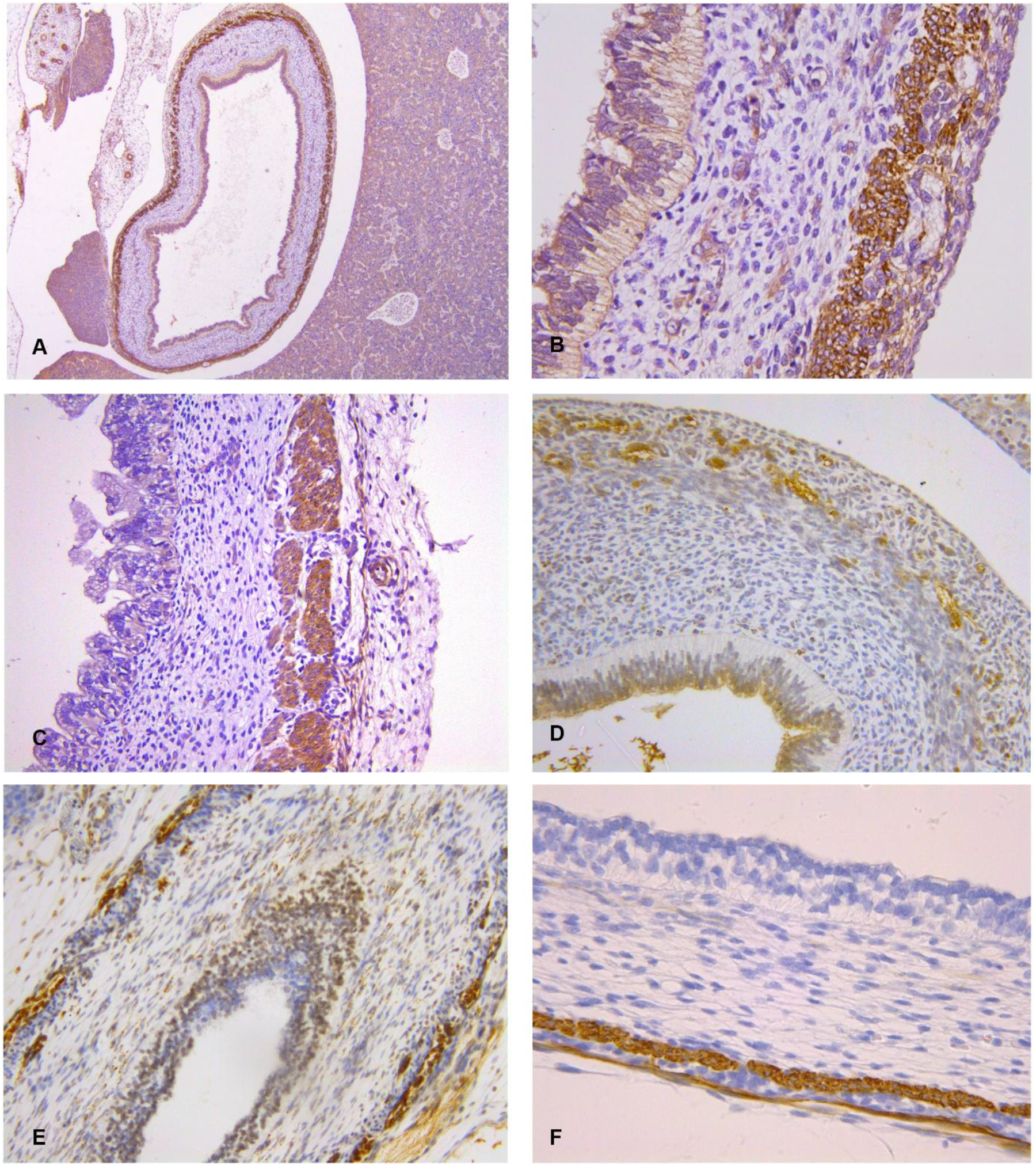
A) ANO1-immunopositivity in inner and outer muscle layer and epithelial cells in stomach,10th week of development, x40; B) ANO1-immunopositivity in stomach in 10th week of development. ANO1-immunopositivity is observed in epithelial cells, endothelial cells of submucosal blood vessels, smooth muscle cells in muscularis propria and elongated cells surrounding the myenteric plexus (corresponding to ICC), x400; C) ANO1-immunopositivity in stomach in 22nd week of development. Epithelial cells show low or absent ANO1 expression. Smooth muscle cells of inner muscle layer and elongated cells around the myenteric plexus (corresponding to ICC) show strong ANO1-positivity, which is absent from smooth muscle cells in outer muscle layer, x250; D) NSE-immunopositivity in ganglionic cells of myenteric plexus in stomach in 8th week of development, x300; E) NSE-immunopositivity in ganglionic cells of myenteric plexus in stomach in 11th week of development, x200; F) Desmin immunopositivity in smooth muscle cells in stomach in 8th week of development, x500.

### The stomach during fetal development

In the 10th to 11th week of development, the wall of the stomach consists of mucosa, submucosa, muscularis propria, and serosa. The epithelium is pseudostratified; however, its height is lower than in the embryonic period and shows signs of differentiation. The epithelial cells display ANO1-positivity, and gastric glands are still not formed (Fig. 2A, 2B). The two muscle layers are discernible in the muscularis mucosa, both exhibiting strong desmin and ANO-1 immunopositivity (Fig. 2B). Elongated ANO1-positive cells entirely surround myenteric plexuses whose cells show strong NSE-positivity, but lack the expression of ANO1 (Fig. 2B, 2E). Within the blood vessels, ANO1-immunopositive endothelial cells are observed.

Between the 14th and 20th week, all layers of the stomach are fully developed. The epithelium is simple columnar, and the gastric glands begin to form. The epithelial ANO-1 immunopositivity is evident, and the pattern of ANO-1 immunopositivity resembles that seen at the start of the fetal developmental period.

In the 25th week of development, epithelial ANO1-immunopositivity is low or absent from cells in surface epithelia and gastric glands. The circular layer exhibits strong ANO1-immunopositivity, whereas in the longitudinal layer, ANO1-immunopositive cells are rare. Very flattened and elongated cells are visible in longitudinal muscle layer and around the myenteric plexuses, which consistently lack ANO-1 immunopositivity. The ANO1-positive cells form a continuous single-cell layer around the margins of the myenteric plexuses. Endothelial positivity is evident in small blood vessels (Fig. 2C).

### Small and large intestine

In the 10th-11th week of development, the primordia of short intestinal villi begin to appear in the small intestine. Intestinal glands are visible in the small intestine; however, they remain absent from the colon (Fig. 3A). The thin, outer longitudinal muscle layer shows strong desmin-positivity and is observable in both the small intestine and the proximal colon (Fig 3B). Where both muscle layers are present, clearly distinctive NSE-positive myenteric plexuses are seen between them (Fig. 1E). The epithelium in both the small and large intestine is pseudostratified, though its height is reduced compared to the 8th week and shows signs of differentiation. Plasmalemmal ANO1-immunopositivity is present across all epithelial cells. The ANO1-positive cells are visible within both the circular and longitudinal layers of the muscularis propria. These cells appear as thin, flattened, elongated structures with bipolar morphology. Flattened, ANO1 strongly positive cells completely enclose the ANO1-negative myenteric plexuses in the form of a continuous belt.

**Figure 3.**
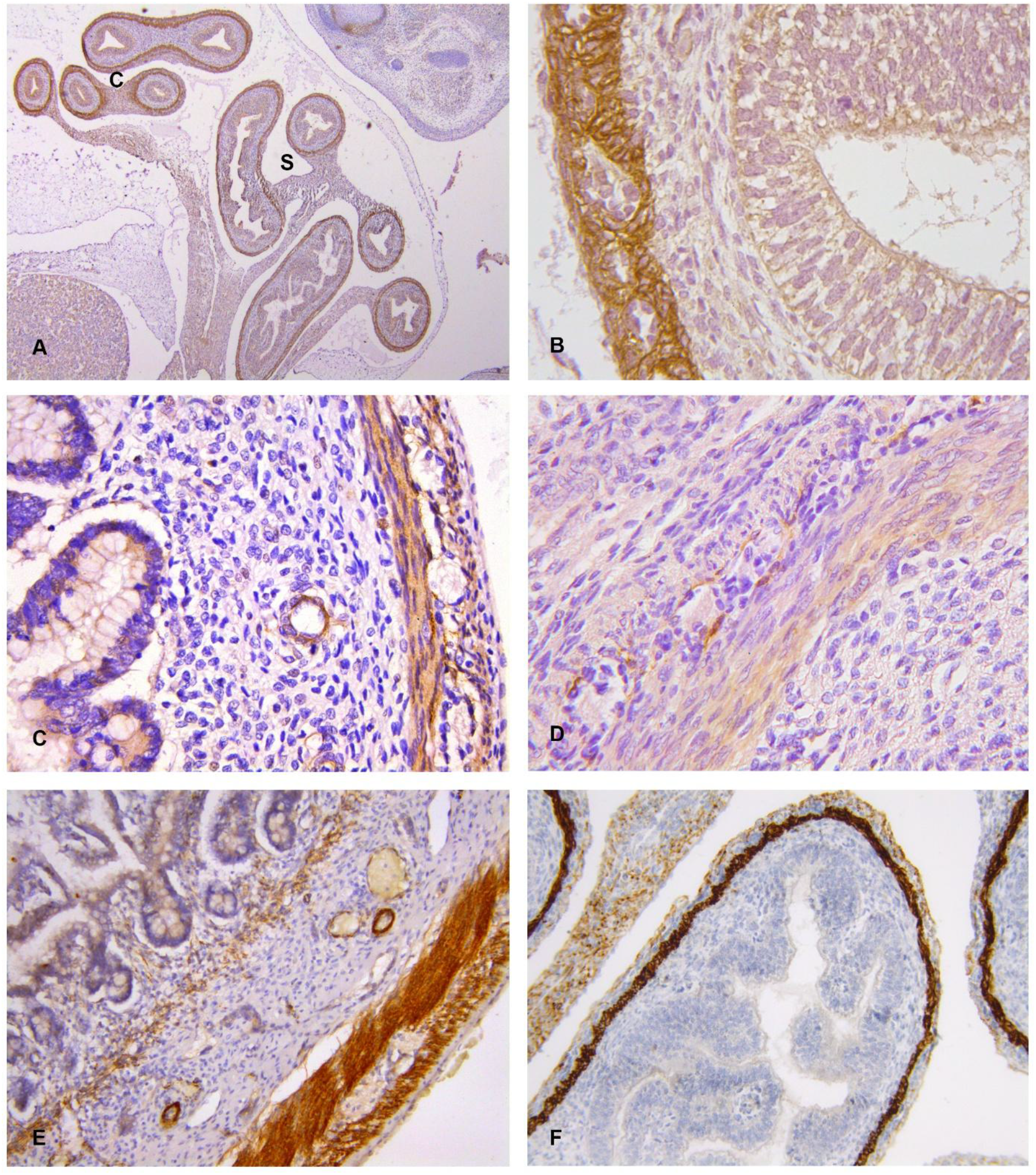
A) Panoramic view of ANO1-immunopositivity in small and large intestine in 10th week of development, x40, S – small intestine, C – large intestine; B) ANO1 immunopositivity in epithelial cells, smooth muscle cells of muscularis propria and cells corresponding to ICC in large intestine in 14th week, x640; C) ANO1 immunopositivity in epithelial cells, smooth muscle cells of muscularis propria and cells corresponding to ICC in large intestine in 17th week, x400; D) ANO1 immunopositivity in smooth muscle cells of inner muscle layer and elongated cells around the myenteric plexus (corresponding to ICC) in large intestine in 19th week, x400; E) ANO1 immunopositivity in epithelial cells, smooth muscle cells of muscularis propria and lamina muscularis musosae, and in cells corresponding to ICC in small intestine in 25th week, x320; F) Desmin-immunopositivity in smooth muscle cells of inner and thin, outer muscle layer in small intestine in 10th week, x200.

Between the 12th and 20th week of development, the intestinal villi are fully formed in the small intestine, and intestinal glands are present in both the small and large intestines. The epithelium consists of simple columnar cells, which exhibit ANO-1 positivity in the apical region and, to a lesser extent, in the lateral areas. In small intestine, ANO-1 immunopositive cells are found within both muscle layers, while in the large intestine ANO1-immunopositivity was low or absent in outer muscle layer. Elongated ANO1-positive cells were seen around the myenteric plexuses whose ganglionic cells lack ANO1 expression. ANO-1-positive cells surrounding the myenteric plexuses display a flattened, elongated shape, and form a complete belt that encircles them. Endothelial cells also show ANO-1 immunopositivity in blood vessels (Fig 3C, 3D).

In the 25th week of development, the small and large intestines are fully developed. The muscularis propria comprises a broader circular layer and a distinctly separate outer muscular layer, both of which stain strongly for desmin. The pattern of ANO1-immunopositivity in the muscularis propria resembles that seen at the 20th week of development. In both the small and large intestines, the lamina muscularis mucosae is clearly visible and contains small, stellate, interconnected ANO1-immunopositive cells. These cells are observed, in continuity with the lamina muscularis mucosae, in the intestinal villi, corresponding to the cells of the muscle of Bricke (Fig. 3E).

## Discussion

Our results show that the stomach, small and large intestine are discernible in histological slides at the end of embryonic development. Their mucosa is lined with pseudostratified epithelium, and the inner circular layer is present in the small and large intestines. In contrast, the thin outer longitudinal muscle layer is observed only in the stomach, where the myenteric plexuses are seen between the two muscle layers. By the 14^th^ week, all histological features of the gastrointestinal tract were developed, including simple columnar epithelium, glands (gastric and intestinal), and a well-defined two-layered muscularis propria (15,16,25,26).

ANO1-positivity in the stomach, small and large intestine was evidenced at the end of the embryonic period in epithelial, endothelial and smooth muscle cells in all examined parts of the gastrointestinal tract. In addition, elongated ANO1-positive cells around the myenteric plexus, corresponding to ICCs were seen around the myenteric plexuses in the stomach. With the maturation and development of the outer muscle layer and myenteric plexus in the small and large intestine, ANO1 showed the same pattern of expression in their ICCs. Ganglionic cells of the myenteric plexus consistently showed a lack of ANO1 expression in both embryonic and later fetal development. Although there are no data concerning ANO1 expression during the development of the gastrointestinal system, the studies report that c-kit, as a widely accepted marker of ICCs, is expressed in the muscle precursors and ICCs in the stomach and the proximal part of the duodenum at the end of embryonic development, suggesting that the differentiation of ICCs and smooth muscle cells begins early during development (26,27,28). This shared positivity likely results from their common mesenchymal origin. Studies showed that these two cell types arise from a common c-kit-positive mesenchymal precursor and that c-kit plays a crucial role in directing their differentiation toward SMC or ICC phenotypes (12,14,29-31). Unlike the ANO1-positivity that is present in all parts of gastrointestinal system at the end of the embryonic development, c-kit immunopositive cells at this period are present only in oesophagus, stomach and proximal part of the duodenum and will appear in the other parts during the 9^th^ and the 10 weeks (16,27,28,32,33). We therefore suggest that ANO1 might be a more specific marker for labelling common mesenchymal progenitors compared to c-kit. We observed ANO1 positivity in SMC and ICC in tissue samples from the 25^th^ week of development, suggesting that these cells continue to express this marker even after the second trimester, unlike c-kit, whose expression on SMC is lost during the early phases of embryonic development. Interestingly, studies on adult human material showed that ANO1 expression in the gastrointestinal tract is strictly limited to ICC and is absent from the smooth muscle cells (10,11,34). This switch in ANO1-positivity might result from the functional maturation of SMCs and the formation of other types of chloride channels on their plasma membrane (35). Available data suggest that the deletion or pharmacological inhibition of ANO1 leads to disorders of gastrointestinal motility, resulting from disorganised and reduced contractility of smooth muscle cells (36-39). Currently, experimental efforts are underway to establish stem cell-based therapies for gastrointestinal disorders; however, most studies focus on differentiating ganglionic enteric cells from embryonic or induced pluripotent stem cells, while attempts to differentiate ICC are scarce (40-42). Dave et al. reported that the transplantation of murine ICC-stem cells into the mice with acute and chronic colitis reduced the severity of symptoms. These cells homed in colon and in vitro showed the ability to suppress T-cell proliferation (43). Given that ANO1 is expressed on ICC and their mesenchymal progenitors, it may be used, in combination with other markers, to establish the specific time points during the differentiation of mesenchymal cells into ICC and SMC, as well as their subsequent maturation.

Endothelial ANO1-positivity was a consistent finding in gastrointestinal tract blood vessels during embryonic and fetal development. Although ANO1 expression has been reported in endothelial cells in the brain, umbilical vein, and heart, we found no data on its expression in endothelial cells during development (44-47). The role of ANO1 in endothelial cells remains incompletely elucidated. Some data suggest that ANO1 promotes vasoconstriction and endothelial dysfunction by generating reactive oxygen species in endothelial cells (44). However, some authors report that activation of ANO1 channels induces vasodilation and a consequent decrease in blood pressure (48,49). According to Garud et al., CaCC activation reduces cytoplasmic chloride ion concentration, thereby activating WNT kinase, which in turn stimulates TRPV4 channels and induces vasodilatation (48).

Our results show that ANO1 is expressed in the epithelial cells in the gastrointestinal tract. This immunopositivity is observed in the pseudostratified epithelium of the embryo and persists, to a lesser extent, in the simple columnar epithelium of fetal samples. As biological membranes, the epithelia play a crucial role in the secretion and absorption of fluids and electrolytes. The expression of ANO1 is reported in intestinal epithelia, where it is assumed to regulate these processes, given the roles of chloride ions to determine the direction of fluid or electrolyte secretion (34,39,50). The lower expression of ANO1 in intestinal epithelium might be explained by the fact that the main anion channel responsible for chloride secretion in these cells is the cystic fibrosis transmembrane conductance regulator (CFTR) (51). Experimental data show an interaction between the CFTR and CaCC signalling pathways, further supported by reports from Benedetto et al. that ANO1 is crucial for the proper membrane function of CFTR (51,52). In addition, ANO1 depletion was shown to reduce calcium-dependent chloride secretion in the small and large intestines of mice, resulting in mild mucosal oedema (50).

In conclusion, ANO1 can be considered as a reliable marker for following the differentiation of SMC and ICC during embryonic and fetal development. ANO1 expression on ICC is pertinent during the fetal development, however the SMC in outer muscle layer of stomach and large intestine loose ANO1-positivity by the 25^th^ week of development. Bearing in mind its early appearance during the embryonic development, ANO1 might be useful for future studies of the differentiation of mesenchymal progenitors into SMC and ICC, as well as their further maturation and in vitro cultivation.

## Acknowledgement

Research reported in this paper was supported by the Project number 451-03-137/2025-03/200113 of the Ministry of Science, Technological Development and Innovations of the Republic of Serbia and the Internal project of the Faculty of Medicine in Niš (INT-MFN no. 38/20). This research was funded by

## Author’s contribution

VP, GR – designing research study; VP – writing the manuscript; GR, AV, MJ – revising the manuscript; VP, GR, JR – analysis of data; DM, VR, BK – performing immunostaining procedures.

## Statement of ethics

The study was reviewed and approved by the Ethics Committee of the University Clinical Center - Niš, number 34794/3, date 1.10.2019.

## Statement of competing interest

The authors declare no conflict of interest.

## Notes

### Competing Interest Statement

The authors have declared no competing interest.

